# The Mouse Cortical Interareal Network Reveals Well Defined Connectivity Profiles and an Ultra Dense Cortical Graph

**DOI:** 10.1101/156976

**Authors:** Răzvan Gămănuţ, Henry Kennedy, Zoltán Toroczkai, David Van Essen, Kenneth Knoblauch, Andreas Burkhalter

**Affiliations:** Univ Lyon, Université Claude Bernard Lyon 1, Inserm, Stem Cell and Brain Research Institute U1208, 69500 Bron, France.; Institute of Neuroscience, State Key Laboratory of Neuroscience, Chinese Academy of Sciences (CAS) Key Laboratory of Primate Neurobiology, CAS, Shanghai 200031, China; Interdisciplinary Center for Network Science and Applications, Department of Physics, University of Notre Dame, Notre Dame, IN 46556, USA.; Department of Neuroscience Washington University School of Medicine, St. Louis, MO 63110-1093, USA

**Keywords:** Neocortex, rodent, tract-tracing, retrograde, lognormal, connectivity, anatomy

## Abstract

The inter-areal wiring pattern of mouse cerebral cortex was analyzed in relation to an accurate parcellation of cortical areas. Twenty-seven retrograde tracer injections were made in 19 areas of a 41 area (plus 7 sub-area) parcellation of the mouse neo-, parahippocampal and perirhinal cortex. Flat mounts of the cortex and multiple histological markers enabled detailed counts of labeled neurons in individual areas. A weight index was determined for each area-to-area pathway based on the Fraction of Extrinsically Labeled Neurons (FLNe). Data analysis allowed cross species comparison with the macaque. Estimation of FLNe statistical variability based on repeat injections revealed high consistency across individuals and justifies using a single injection per area to characterize connectivity. The observed lognormal distribution of connections to each cortical area spanned 5 orders of magnitude and revealed a distinct connectivity profile for each area, analogous to that observed in macaque. The resulting graph has a density of 97% (i.e. 97% of connections that can exist do exist), considerably higher than the 66% density reported for the macaque. Our results provide more sharply defined connectivity profiles and a markedly higher graph density than shown in a recent probabilistic mouse connectome.

## Introduction

The concept of the cortical area is rooted in the notion of localization of function in the cortex, where individual areas are posited to have a distinct architecture, connectivity, function, and/or topographic organization (Felleman and Van Essen, 1991). The mouse is increasingly used as a model system for investigating the cortex, where complex sensory (Ferezou et al., 2007), motor (Li et al., 2016) and cognitive (Carandini and Churchland, 2013; Kim et al., 2016; Manita et al., 2015) functions have been shown to depend on interactions among cortical areas via inter-areal connections, as well as on dynamic control involving higher-order thalamic nuclei (Mease et al., 2016; Sherman, 2016). The highly interactive nature of cortical processing has motivated recent efforts to investigate the statistical properties of inter-areal networks and the development of large-scale models of the cortex that may provide insights into brain function in health and disease (Bullmore and Sporns, 2012).

Our understanding of distributed hierarchical processing within the brain has benefited from investigations of the cortical network at the systems level using tract tracing. Felleman and Van Essen (1991) relied on collated data from numerous studies using diverse methods and approaches. However, the low sensitivity of early pathway-tracing methods and differences in experimental procedures across labs limited the reliability of some of the statistical features of large-scale cortical networks that can be derived from this database (Kennedy et al., 2013). Most notable is the underestimation of the density of the cortical graph (i.e. the fraction of connections that can exist which do exist). This in turn narrows the range of plausible models of cortical networks (Markov et al., 2013b). These considerations have motivated sensitive tract-tracing methods and consistent areal parcellation (Bassett and Bullmore, 2016). The high density and wide range of connection strengths of the cortical graph (Markov et al., 2014; Markov et al., 2011) makes it particularly important to quantify the *weights* of the connections linking different cortical areas (Ercsey-Ravasz et al., 2013; Oh et al., 2014; Ypma and Bullmore, 2016), and to characterize the variability of connection strengths across repeat injections.

Studies of the cortical anatomy have systematically failed to report quantitative connectivity data, in part due to earlier claims that connection weights in cortical pathways are highly variable with a 100-fold weight range (MacNeil et al., 1997; Musil and Olson, 1988a, b; Olson and Musil, 1992; Scannell et al., 2000). These reports suggesting a high variability in connection weight could raise serious problems concerning the anatomical specificity of the cortical network, particularly in high density graphs, and would appear to argue against the concept of each area having a distinctive fingerprint or connectivity profile defining its functional specificity (Bressler and Menon, 2010; Markov et al., 2011; Passingham et al., 2002). More recently, the specificity of connection weight has been reexamined in macaque. In particular, retrograde tracer studies showed that each target area receives inputs from a specific set of areas, each individual projection displaying a characteristic weight (Markov et al., 2014; Markov et al., 2011). These studies compared connection strengths across individuals using consistent criteria showed that in macaque the weights range over 5-orders of magnitude, several orders of magnitude larger than the variability observed from a single projection. For a given target area the set of areas with their specific weights constituted a characteristic connectivity profile. Hence, in addition to having weights, it is imperative that the mouse data base is investigated for its statistical variance in order to estimate its consistency.

A recent parcellation using multiple modalities indicates that the mouse cortex contains about 40 distinct areas (Wang et al., 2012), significantly fewer than reported for humans (Glasser et al., 2016) and the macaque monkey (Van Essen et al., 2012). The weight-distance relations observed in macaque cortical connectivity data lead to a one parameter predictive model that captures multiple features of the cortical network including its spatial embedding, wire minimization, frequency distribution of motifs, global and local efficiencies and a core-periphery architecture (Ercsey-Ravasz et al., 2013). Spatial embedding constrains numerous geometrical features in a similar fashion in both macaque and mouse cortex (Horvat et al., 2016).

A recent mouse connectome study at the mesoscale level used anterograde tracer injections to obtain brain-wide weighted data (Oh et al., 2014). Because most of the reported injection sites (>70%) spanned multiple areas, connectivity at the level of individual areas was inferred using a computational model involving several theoretical assumptions. The resultant probabilistic model of the mouse cortical connectome generated a density for the inter-areal graph of 35-53%, much lower than the 66% reported for the macaque cortical graph (Markov et al., 2014). The lower density reported in the mouse is surprising given the expected increase in density with decreasing brain size (Horvat et al., 2016; Ringo, 1991). Furthermore, an earlier tracer study of mouse visual cortex (Wang et al., 2012) reported a higher subgraph density (99%) than the 77% found for visual areas in the Oh et al., 2014 study. These results suggest that the computational procedure used in the Oh et al., 2014 study to infer the connectivity of single areas from injections involving multiple areas might have resulted in significant numbers of false negatives. Ypma and Bullmore reanalyzed the Oh et al., 2014 dataset and estimated a whole-cortex graph density of 73% (Ypma and Bullmore, 2016). Although a computational modeling approach to connectomics deserves future exploration, in the present study we focused on an empirical approach that is essentially deterministic insofar as it depends on direct anatomical observations.

We investigated mouse cortico-cortical connectivity and addressed two key issues: the density of the mouse cortical graph and the consistency of connectivity profiles. We minimized experimental variability by targeting injections of a retrograde tracer in post-hoc identified areas rather than anterograde tracing on a fixed grid of injections (Oh et al., 2014). Our choice of retrograde tracer provides several advantages for quantifying the strength of connections (Markov et al., 2014). We coupled retrograde tracing with flatmounting the cortex, which is advantageous when combined with multiple histological stains that reveal high-resolution areal maps (Qi and Kaas, 2004; Sincich et al., 2003; Wang and Burkhalter, 2007; Wang et al., 2011; Wang et al., 2012). Our experimental approach constitutes a positive identification of injected cortical areas and of areas where the retrogradely labeled neurons are located.

We show that modeling of the variability observed following repeated tracer injections displays statistical consistency, similar to that obtained in macaque (Markov et al., 2014). Our results show that the mouse cortex is ultra-dense with a graph density of 97%, significantly higher than the probabilistically-based range of 35-73% (Oh et al., 2014; Ypma and Bullmore, 2016). The high density of the mouse cortical graph suggests that the activity pattern of a given area is interrelated via its connectivity profile to a widespread pattern of influences across the cortex. While reduction in brain size is expected to entail an increase in graph density, it would seem unlikely that this would be accompanied by a level of variability of weights as reported in Oh et al., 2014 which in conjunction with the high graph density would entail a much reduced specificity of the structure and function of the cortical network compared to macaque. Our analysis of the present retrograde labeling revealed variance of connectivity and connectivity profiles in the mouse comparable to those observed in macaque (Markov et al., 2014). Likewise, our deterministic approach leads to a weight-distance relationship that is quantitatively similar in mouse and macaque (Horvat et al., 2016), which is not the case using the modeled data from the Oh, et al., 2014 study. Our findings provide important structural underpinnings of inter-areal processing in the mouse, which is increasingly used for the investigation of cognitive, sensory, motor and behavioral functions.

## MATERIALS AND METHODS

Experiments were performed in 102 male and female C57BL/6J and PV-Cre (Hippenmeyer et al., 2005)(Jax: 008069), x Ai9 reporter mice (Jax: 007905), harboring the loxP-flanked STOP cassette, which prevented the transcription of the tdTomato protein driven by the chicken β-actin (CAG) promoter (Madisen et al., 2010). The Cre-mediated recombination, resulted in expression of the red fluorescent tdTomato (tdT) protein in parvalbumin (PV)-positive GABAergic neurons. All experimental procedures were approved by the institutional Animal Care and Use Committee at Washington University.

### Tracer injections

For tracer injections, mice were anesthetized with a mixture of Ketamine (86 mg · kg -1) and Xylazine (13 mg · kg-1, i.p) and secured in a headholder. Body temperature was maintained at 37°C. Left-hemisphere tracer injections were made by inserting a glass pipette (20 μm tip diameter) through the dura into the brain and injecting the retrograde tracer Diamidino Yellow (DY; 50 nl, 2% in H2O; EMS-Chemie, Gross-Umstadt, Germany) by pressure (Picospritzer, Parker-Hannafin). Injections were aimed stereotaxically 0.35 mm below the pial surface and often required pulling back the pipette to correct proportionally for dimpling of the dura. Cases in which DY spilled into the white matter were excluded from the analysis. The origin of the coordinate system was the intersection between the midline and a perpendicular line drawn from the anterior border of the transverse sinus at the posterior pole of the occipital cortex. The injection sites, identified as 120-280 μm-wide crystalline deposits of DY (Figure S2A, D, F) in 19 parcels (18 areas of which SSp was injected in two subareas) of occipital, temporal, insular, parietal, restrosplenial, motor, cingulate and prefrontal cortex (anterior/lateral in mm): ACAd (6/0.1), AL (2.4/3.7), AM (3/1.7), AUDpo (2.3/5), DP (4.5/2), GU (6/4), LM (1.4/4.2, MM (1.9/1.6), MOp (6.45/2), P (0.7/4.1), PL (6.8/0.1), PM (1.9/1.6), RL (2.8/3.3), RSPd (2.1/0.4), SSp-bfd (3.4/3.25), SSp-un (3.4/2.4), SSs (3.75/3.25), V1 (1.1/2.4-2.9), VISC (4.2/4.5). In all cases, DY deposits were confined to individual areas (Figure S2A-F). Note that under fluorescence illumination, injection sites appeared larger, but nevertheless showed no apparent spread to neighboring areas.

### Histology

Four days after tracer injection, mice were deeply anesthetized with an overdose of Ketamine/Xylazine and perfused through the heart with phosphate buffered saline, followed by 1% paraformaldehyde (PFA) in 0.1M phosphate buffer (PB, pH 7.4). The cortex was immediately separated from the rest of the brain. To unfold the cortex the tissue was placed on a glass surface, pial surface down. Using microsurgical knives, the hippocampus was separated from the cortex by detaching the alveus from the external capsule and folding it back with the dorsal side up, but still attached by the ventral subiculum to the cortical amygdala and entorhinal cortex. A small incision was made to separate medial from lateral orbital cortex. Working in a posterior direction the white matter was split between the corpus callosum and the cingulate bundle, enabling the unfolding of the medial wall containing medial orbital, prefrontal, cingulate, and retrosplenial cortex. The tissue was then transferred white matter down onto a filter paper placed on a sponge sheet and covered with a glass slide (25x75x1 mm). The assembly was postfixed in a petri dish filled with 4% PFA and stored overnight at 4°C. After postfixation the tissue was cryoprotected in 30% sucrose and 40 μm thick sections were cut on a freezing microtome in the tangential plane.

In order to assign in each mouse the injection site and DY labeled neurons to specific cortical areas we developed a parcellation scheme based on the distinctive distribution of PVtdT expression. This approach allows charting the location of neurons in sections that are accurately parcellated. It eliminates staining for areal markers, and avoids loss of signal and the associated risk of secondary labeling by leakage from retrogradely DY labeled cells. To determine whether PVtdT density reliably labels distinct parcels, we looked for correlations between PVtdT borders with those revealed by the expression of M2 muscarinic acetylcholine receptor (M2), vesicular glutamate transporter 2 (VGluT2) and cytochrome oxidase (CO) reactivity. All of these markers were employed previously to segment rodent cortex (Ichinohe et al., 2003; Wang et al, 2011; Wang et al, 2012). In order to compare all four patterns we stained alternate series of tangential sections from flatmounted PVtdT-expressing cortex with an antibody against M2 (MAB367, Millipore) or VGluT2 (AB2251, Millipore) and used fluorescent secondary antibodies for visualization. Alternatively, we used non-fluorescent immunohistochemical ABC staining methods to visualize M2 and VGluT2 expression and histochemistry to reveal CO reactivity. In each case, alternate sections were stained for Nissl substance to reveal the cytoarchitectonic landmarks annotated in the Allen Brain Atlas. The expression patterns were imaged under a microscope equipped for brightfield and fluorescence illumination.

For plotting DY labeled neurons, the sections were mounted onto glass slides and analyzed under UV fluorescence (excitation: 355-425 nm, emission: 470 nm) at 20×, with a microscope controlled through a computer with the Mercator software package, running on ExploraNova technology. Labeled neurons were contained in all 12–24 sections per hemisphere. Digital charts of the coordinates of DY labeled neurons across each section were stored in the computer. When charting was complete, the sections were imaged for PVtdT (excitation 520-600 nm, emission 570-720 nm), then stained for Nissl substance with cresyl violet and imaged under bright field illumination. The images of the sections were acquired using MorphoStrider software (ExploraNova).

### Segmentation

The digitized charts of the neurons and the images of the corresponding sections were brought to a common scale and aligned in Adobe Illustrator. The sets of images from each brain were segmented using the regional patterns in the densities of PVtdT expression and, where helpful, of Nissl stained cell bodies. (For each case, two investigators did mutually complementary work in order to minimize the errors from manual drawing). The assignment of the labeled neurons to their respective cortical areas resulting from the segmentation was done with in-house software, written in Python 2.7. The fraction of labeled neurons per area (FLNe) was estimated as the number of labeled neurons expressed as a fraction of the total number of labeled neurons in the cortical hemisphere extrinsic to the injected area.

## Results

We first describe the experimental techniques used to parcellate the cortex and to determine the numbers of labeled neurons in identified cortical areas. We then describe the statistical variation of connectivity across individuals in order to compare the consistency and reliability of the connectivity profile for the retrograde tracer data generated in the present study with an earlier study in macaque (Markov et al., 2014; Markov et al., 2011) and the anterograde tracing data of Oh et al., 2014. Finally, we compare the graph density obtained with the retrograde tracer data and detail the different densities obtained in the Oh et al., 2014 study.

### High-resolution cortical parcellation

Given the small mouse cortex, the spatial precision by which tracer injections can be mapped onto the underlying structure is vital to success. To this end we used PVtdT mice, which allowed us to segment the cortex into 24 parcels excluding entorhinal, hippocampal and piriform cortex (Figure 1A) and provided stable landmarks for the identification of 41 distinct areas (Figure S1B) in injected brains. This allowed mapping the sources and targets of connections and avoided *post hoc* staining for areal markers, which interferes with DY-labeling, obviate aligning parallel sections and avoid assigning labeled cells to a standard template. This approach accounts for the variability of the parcellation across individuals (Krubitzer and Seelke, 2012) and therefore significantly differs from that employed in two recent studies of the mouse connectome (Oh et al., 2014; Zingg et al., 2014), which mapped corticocortical projections onto a standard Allen Reference Atlas (ARA, (Dong, 2008)), generated by averaging variations of background fluorescence across hundreds of cortices. This Common Coordinate Framework (CCF, Allen Institute, brain-map.org) has become a widely used parcellation of mouse cerebral cortex (Figure S1A). Our choice of using PVtdT expression for areal identification was inspired by the work of Saleem and Logothetis ((Saleem, 2012)), who used PV immunostaining to delineate many cortical areas in rhesus monkey. There is no *a priori* reason to assume that the expression pattern of PVtdT outlines areal boundaries. As this is of course true for any singular architectural measure, we compared the pattern of PVtdT expression with the patterns of immunolabeling for M2, VGluT2, histochemical reactivity for CO and Nissl substance. Based on previous observations (Wang et al., 2011) our expectation was that comparing different markers would reveal overlapping or complementary spatial expression gradients reflecting areal borders across individuals.

**Figure 1.**
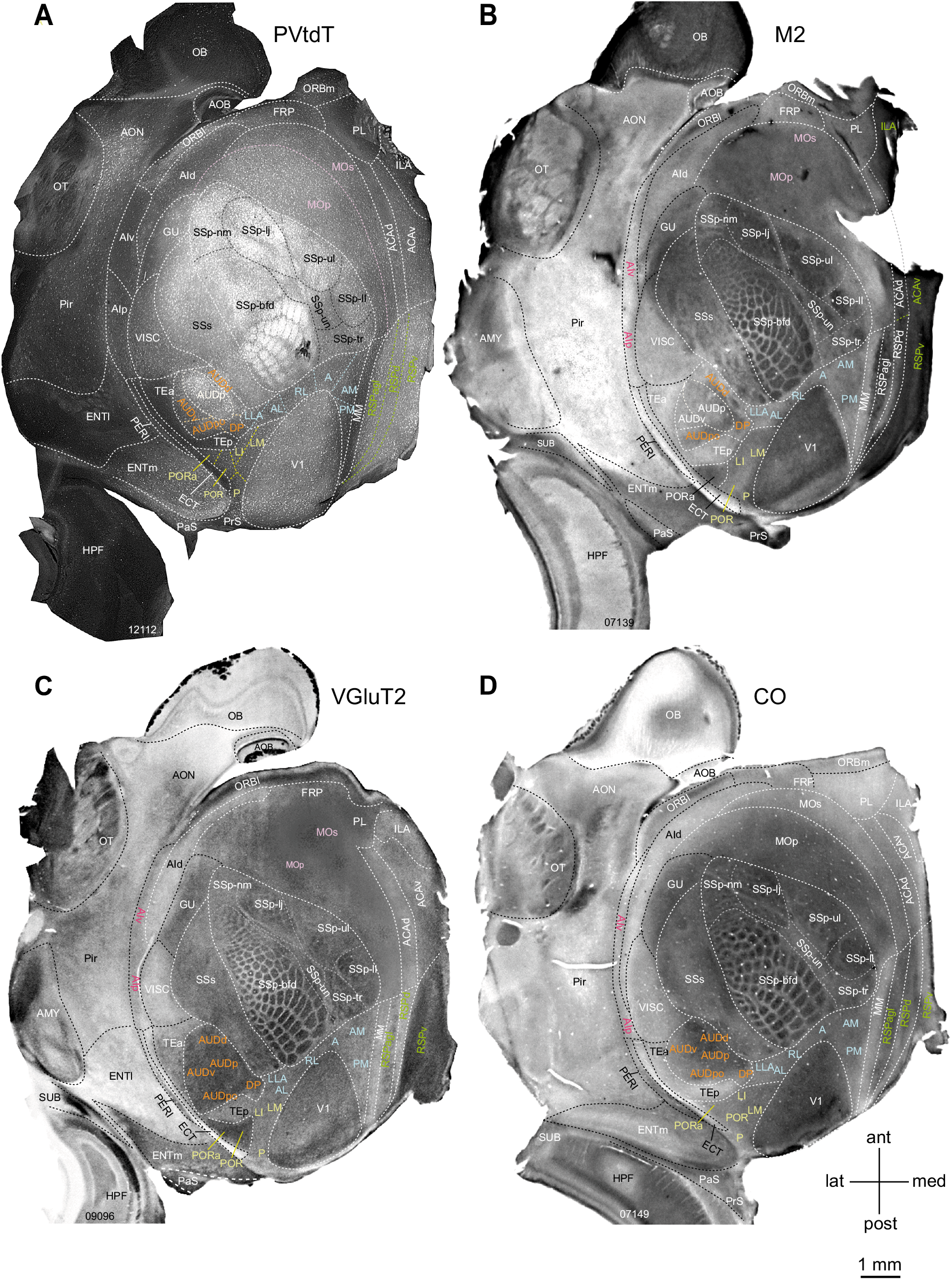
Expression of different markers in layer 3/4 of flatmounted left mouse cerebral cortex. (**A**) Tangential section showing tdTomato fluorescence in PV-containing interneurons (bright white labeling). Parcels outlined by white dashed lines and labeled by black and white letters were positively identified by PVtdT expression. Black dashed lines indicate subdivisions within primary somatosensory (SSp) cortex representing different body parts. Colored letters denote known areas contained within distinct compound parcels (orange, yellow, blue, green, pink) in which PVtdT-expression exhibits similar intensity and reveals no detectable subdivisions. Colored dashed lines indicate presumptive borders between these areas. (**B**) Bright field image of tangential section stained with an antibody against the M2 muscarinic acetylcholine receptor (dark staining). Areas outlined with white and black dashed lines and denoted with white and black letters were positively identified as distinct parcels. Areas denoted in orange, yellow, blue, green, pink and red letters indicate known areas contained within distinct, but uniformly M2-labeled parcels. (**C**) Bright field image of tangential section stained with an antibody against VGluT2 (dark staining). Areas outlined with white and black dashed lines and denoted in white and black letters were positively identified as distinct parcels. Areas denoted in orange yellow, blue, green pink and red letters indicate known areas contained within distinct, but uniformly VGluT2-labeled parcels. (**D**) Bright field image of tangential section reacted for cytochrome oxidase (CO) activity (dark staining). Areas outlined with white and black dashed lines and denoted in white and black letters were positively identified as distinct parcels. Areas denoted in orange, yellow, blue, green and red letters indicate known areas contained within distinct, but uniformly COlabeled parcels.

By comparing PVtdT, M2, VGluT2 and CO expression patterns, we positively identified in PVtdT maps 24 parcels (one of which was subdivided into 7 subareas). Ten additional areas in M2, VGluT2 and CO maps (POR, PORa [previously referred to as 36p, (Wang et al., 2011)], AUDv, AUDpo, AUDv, DP, MOs, RSPagl, RSPd, RSPv) were identified with respect to the PVtdT map in each DY-injected mouse by their location relative to neighboring areas. The complete borders of 7 known areas (P, LI, LLA, RL, A, AM, PM; (Wang et al., 2007)) could not be positively identified by any of the markers we used but were nevertheless inferred by their stereotypical size, shape and location relative to PVtdT landmarks. The grand total was 41 areas, of which SSp was subdivided into 7 subareas (Figure S1B). We estimate that areal boundaries were drawn in most regions with < 150 μm accuracy. For additional information on parcellation see Supplementary Information.

### Retrograde tracing with DY

Of a total of 102 DY injections, 27 injection sites were confined to 17 distinct areas and two subareas of SSp. Typical examples of such injection sites are shown in Figures S2A-F. These included repeat injections of AM (n=2), LM (n=3) (Figure S2D), RL (n=2), SSp-bfd (n=2), V1 (n=4) (Figure S2A-C) and singular injections of ACAd (n=1), AL (n=1) (Figure S2E), AUDpo (n=1), DP (n=1), GU (n=1), MM (n=1), MOp (n=1), P (n=1), PL (n=1), PM (n=1) (Figure S2F), RSPd (n=1), SSp-un (n=1), SSs(n=1), VISC (n=1). In all cases, retrograde DY labeling was exceptionally bright and labeled large numbers of neurons (nuclei and cytoplasm) in multiple areas distributed across the cortex.

Representative examples of the DY labeling at low power are shown in area V1 (Figure S2B,C). Note, cell counts were obtained from charts at higher magnification than those shown in Figure S2B, C. The area V1 injection was confined to the lower peripheral visual field representation near the tip of V1 (Marshel et al., 2011). As expected from previous axonal tracing and topographic mapping experiments (Garrett et al., 2014; Marshel et al., 2011; Wang et al., 2007), retrogradely DY labeled neurons were clustered at the junction of areas LM, AL, LI and LLA. Additional clusters of labeled neurons were found at retinotopically corresponding locations within RL, A, AM, PM, MM, P, POR, PORa. In temporal cortex, DY labeling was contained in most areas of auditory cortex (AUDp, AUDPo, AUDd and DP) and the ventral portion of the posterior temporal area (TEp). On the medial wall, DY labeling was clustered in dorsal retrosplenial cortex (RSPd), the secondary motor cortex (MOs), the dorsal and ventral anterior cingulate areas (ACAd, ACAv) and in the prelimbic (PL) and infralimbic (ILA) areas. At the rostral end of cortex, labeling in orbitofrontal cortex was confined to ORBl. In transitional cortex, DY labeled cells were found in posterior ectorhinal (ECT) and the medial entorhinal cortex (ENTm). A high density of DY-labeled cells was found throughout the dorsal and ventral claustrum.

The utility of the parcelation scheme and the correct allocation of labeled neurons to individual cortical areas is upheld by the spatial maps of labeled neurons such as shown in Figure 2, which reveals a pronounced modulation of density with respect to area borders.

**Figure 2.**
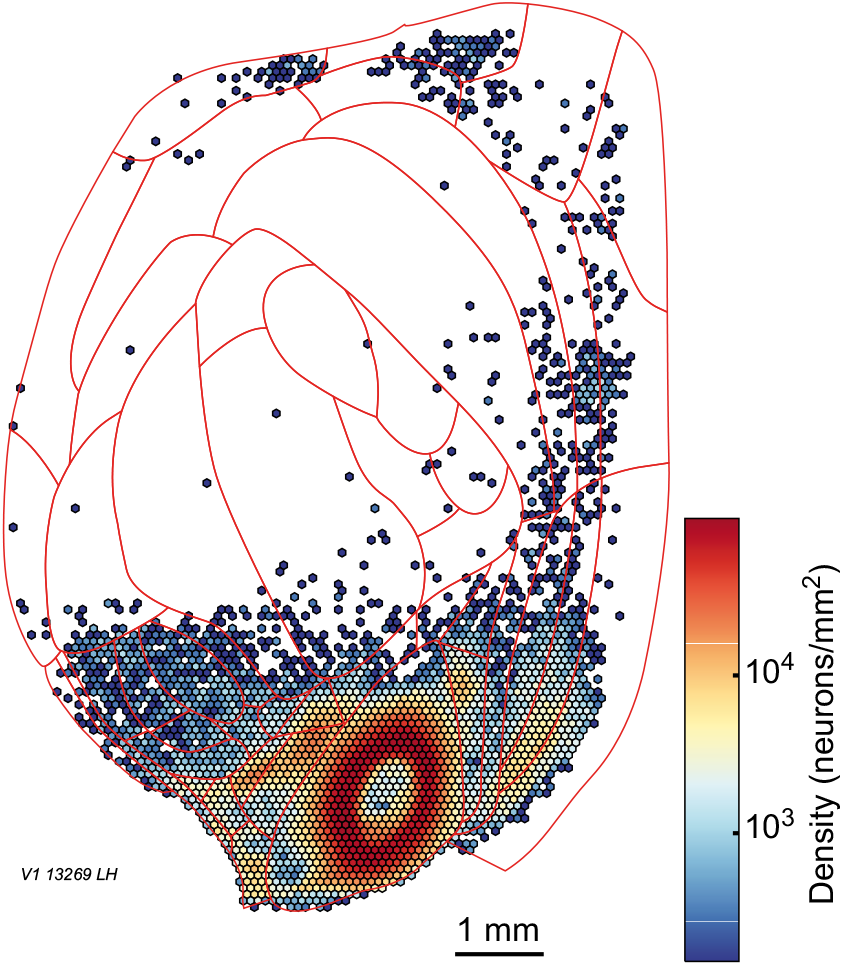
Two dimensional spatial distribution with respect to areal borders of retrogradely labeled neuron following an injection in area V1.

### Weight overdispersion

Quantifying variance allows accurate statistical inferences based on the data and also estimation of the uncertainties associated with observed weight values. This allows evaluation of connectivity profiles and provides useful constraints on how well single injections can be used to estimate connectivity profiles (Markov et al., 2014; Markov et al., 2011). Variability in the data was modeled by analyzing the statistical properties of connections resulting from repeat injections of tracers (Figure 3 A-B). The connection strengths are based on the counts of the number of labeled neurons in each parcellated source area following an injection of DY in a given target area. Since the counts for each source area were normalized by the total number of neurons projecting to the target area, minus neurons labeled within the injected target area, the FLNe provides a measure of the relative strength of each projection. While the normalization constrains the FLNe values to be in the interval (0, 1), they are still based on counts and display properties of discrete data. In statistical models of count data, the variance typically depends upon the mean. An important question is whether the variance is overdispersed with respect to the mean. For additional information on overdispersion see Supplementary Information. In the negative binomial family, the variance depends on the mean via a dispersion parameter. A guide to what that parameter should be is obtained by examining plots of the square root of the variance or standard deviation (SD) of FLNe against the mean of FLNe for repeated injections in the same cortical area. Figure 3A, B shows plots of the SD versus the mean for multiple injections in two cortical areas in which the log(SD) increases approximately linearly with the log(FLNe).

**Figure 3.**
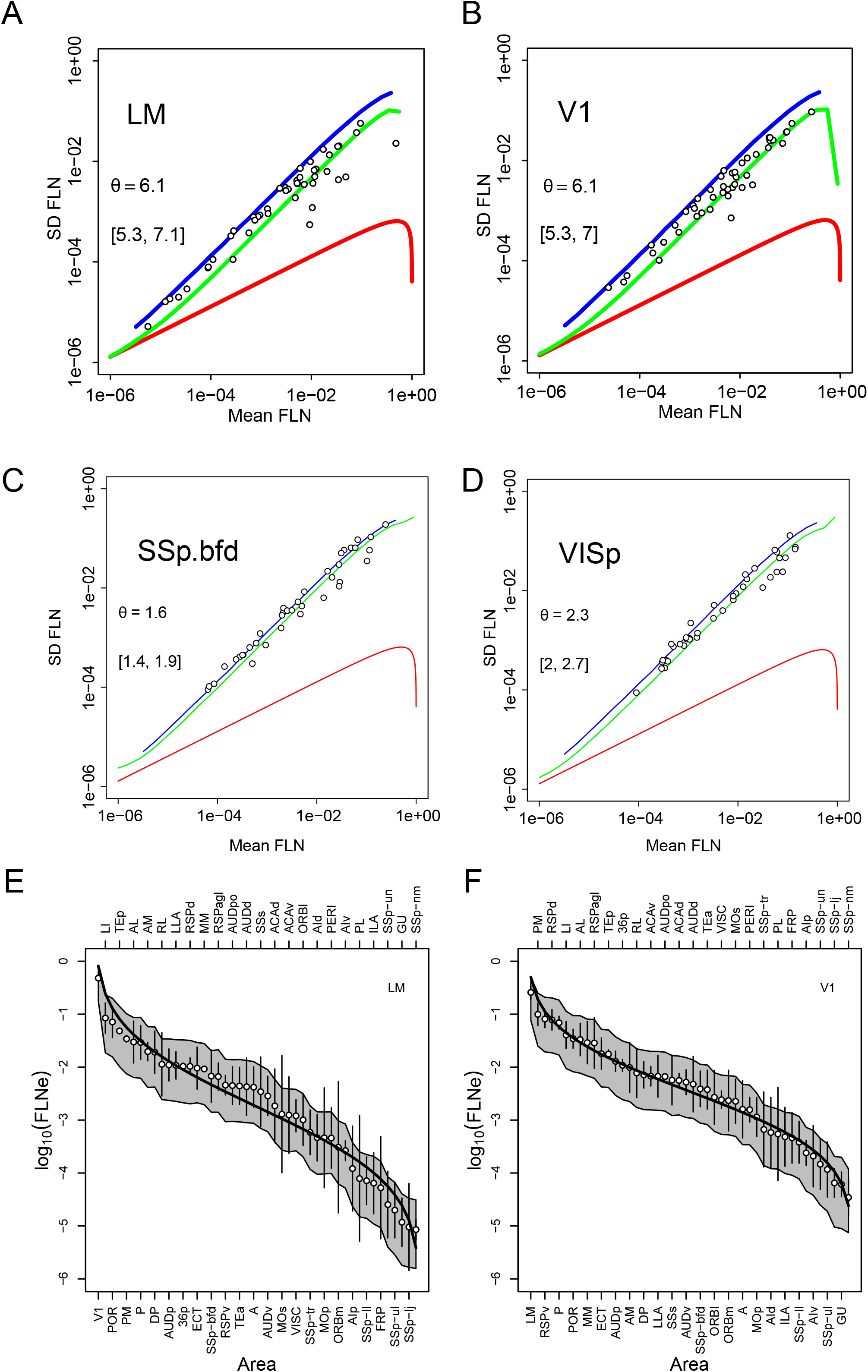
Variance and lognormal distribution of FLNe. **A-D** Repeat injections SD as a function of the mean; θ, dispersion parameter, Red curves, Poisson distribution; blue, geometrical distribution; green, negative binomial; brackets, 95% confidence interval. (**A-B**) retrograde tracers (present study) reveal a negative binomial distribution in LM (**A**), and V1 (**B**). (**C-D**) anterograde tracer data in Oh et al., 2014 reveal a geometric distribution. **E-F**: Lognormal distribution of retrograde tracing data in present study, observed means (white dots) ordered by magnitude, SEMs (error bars) of logarithms of the FLNe for the cortical areas projecting on the injected area. V1 (n=4), LM (n=3). Black curves, the expected lognormal distribution for an ordered set of projections of size n, equal to the number of source areas. The grey envelopes around each curve indicate the 95% confidence intervals obtained by simulating 10000 sets of count experiments drawn from a negative binomial distribution, with means of counts and dispersion parameter as the data.

Both the Poisson (red curves in Figure 3A, B) and the geometric (blue) distributions are extreme cases of the negative binomial distribution family that is frequently used to model overdispersed count data (Hilbe, 2007; Lindsey, 1999; Venables and Ripley, 2002). As explained in the Supplementary information, this is obtained from a mixture of Poisson and gamma distributions and yields a distribution that depends on two parameters, the mean and the dispersion, *θ*. We performed simulations of data sets (described in Supplementary data) for a wide range of dispersion values to estimate the expected relation between SD and mean FLNe as a function of the dispersion. We then compared these curves with the values in Figure 3 and estimated a dispersion value that minimized the error between the simulation and the data, which constitutes the green curves for each plot with the dispersion parameter and the 95% confidence interval (indicated in brackets).

The dispersion values for the retrograde tracer data from the present study are shown in Figure 3A, B. Overall, the results indicate a negative binomial distribution with dispersion values of 6, which provides a reasonable estimate of the expected variability for neural counts obtained with retrograde tracing data in mouse. The *θ* estimate of 6.1 for V1 and LM is somewhat smaller than that obtained in the macaque, 7.9 (Markov et al., 2011), suggesting marginally greater levels of overdispersion across animals in the mouse data sets. Likelihood ratio tests performed on the data reject dispersion values as low as 1 (blue curves) (see Supplementary information) which would correspond to a heavy-tailed geometric distribution. These findings demonstrate that overdispersion is a systematic phenomenon of neural retrograde count data in both macaque and mouse. Thus, overdispersion needs to be taken into account in statistical evaluation of such data since ignoring it would lead to anti-conservative estimates of the probabilities of significant differences in connection strengths, i.e. erroneously assigning significance to small differences.

Repeat injections make it possible to examine variability in the raw data of Oh et al., 2014 (Figure 3C, D). Although not based directly on counts, we can apply the same framework to examine the variability in relation to the mean. In contrast to the retrograde tracer data, the anterograde data of Oh et al., 2014 show a dispersion of 1.6 for the somatosensory barrel field and 2.3 for area V1 and therefore in both cases indicates a much more overdispersed data set compared to the retrograde tracing data in the present study.

### Lognormal distribution of weights

An interesting aspect of cortical organization is that physiological and anatomical features ranging from synaptic densities to areal distances are distributed according to a lognormal distribution (Buzsaki and Mizuseki, 2014). This is a skewed, heavy-tailed distribution typically characterized by relatively few very strong and very weak values with many intermediate ones. A lognormal distribution as a characteristic of the distribution of input strengths to a cortical area was first demonstrated for the macaque (Markov et al., 2011) and subsequently reported in mouse (Oh et al., 2014; Wang et al., 2012).

Figure 3E, F shows the ordered log (FLNe) values (points) for area V1 and LM, the areas analyzed in Figure 3A, B. The range of values spans 4 to 5 orders of magnitude. The error bars are twice the standard errors of the means. The solid curves are the predicted values for ordered Gaussian variables with the same mean and SD as the data set. To evaluate whether differences were significant, we simulated 10000 count data sets from a negative binomial distribution with the same means as the data and with the dispersion parameters obtained from the analyses displayed in Figure 3A, B. For each data set, we normalized the counts by the total to obtain simulated FLNe values. From these distributions, we estimated the 2.5% and 97.5% quantiles to obtain a confidence interval. The grey envelopes in each plot indicate the region defined by these confidence intervals. In general, the ordered means fall within the grey envelopes, indicating that the differences from the lognormal prediction are not significant.

### Consistency of weak projections

The repeatability of connections following repeated injections in a given area across individuals makes it possible to evaluate the consistency of individual pathways. By consistency, we refer specifically to whether a connection is systematically present across injections. In macaque, Markov et al. (Markov et al., 2014; Markov et al., 2011) reported a few cases in which observed projections were inconsistent, i.e., no neurons were observed projecting from a particular source area to a target in at least one of the repeat injections. For discrete distributions, it is expected that there is a non-zero probability of observing no neurons, even if the projection exists, simply based on sampling variability. The probability of observing such cases would be expected to increase as the mean size of the projection decreased. In fact, the probability distributions predict the expected incidence of such missing connections. For Poisson distributed data, the probability of observing zero neurons is *e^−μ^*, where *μ* is the mean number of neurons in the projection. For a negative binomial distribution, the probability of observing zero neurons is 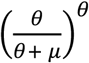.

Deviations from such predictions in the direction of too many observations of zero labeled neurons (leading to zero-inflation) or too few (leading to zero-deflation) constitute evidence against the statistical model used for the data. An alternative is to explain such deviations as evidence that the connections are actually absent in some animals, providing evidence of individual differences in the connectivity pattern. The overdispersion of the count data might be taken as evidence of individual differences in connectivity strength that could be due to genetic or environmental influences. However, without detailed knowledge of what factor(s) are responsible for the overdispersion, such a hypothesis cannot be confirmed.

Figure 4A displays mouse data from 13 repeat injections in target areas (2xAM, 3xLM, 2xRL, 2xSSp-bfd, 4xV1) where the mean number of neurons for a given projection across the multiple injections is plotted as a function of the total number of neurons counted for each injection. Thus, points are individual projections. Those resulting from different injections but for the same target areas are at identical ordinate values, as they all have the same mean and different abscissa values as injections differ only in the total number of neurons labeled in the brain. When no neurons from a given source area were observed in an injection but were observed in at least another repeat injection, the point is plotted as a white disk but is black otherwise. Of the 598 possible connections, 97.1% were present and 17 were absent. Absent connections are concentrated in the lower half of the diagram where low mean number of neurons per projection are found. We first analyzed this by determining whether the features log(Mean) and log(Total) could be used to linearly classify the presence and absence of connections. This can be implemented as a logistic regression in which the expected value of the binary variable (Presence/Absence) is predicted by the two features. The continuous line in Figure 4A indicates the estimated linear classifier for which the probability of a connection being present is 0.95 for this model. Its negative slope suggests a dependence of connectivity on both features such that small injections would lead to higher probabilities of absence at high mean connection strengths. However, only the mean of projection was found to contribute significantly to the linear classifier, (log(Mean): z = 5.57, p = 2.56*10^−8^; log(Total): z = 1.7; p = 9.6 10^−2^), thereby rejecting this hypothesis. The 95% classifier based only on the log(Mean) is indicated by the dashed line and corresponds to a value of 24 neurons. We expect, therefore, that projections revealed in an injection, which are stronger than 24 neurons, to be highly consistent across individuals.

**Figure 4.**
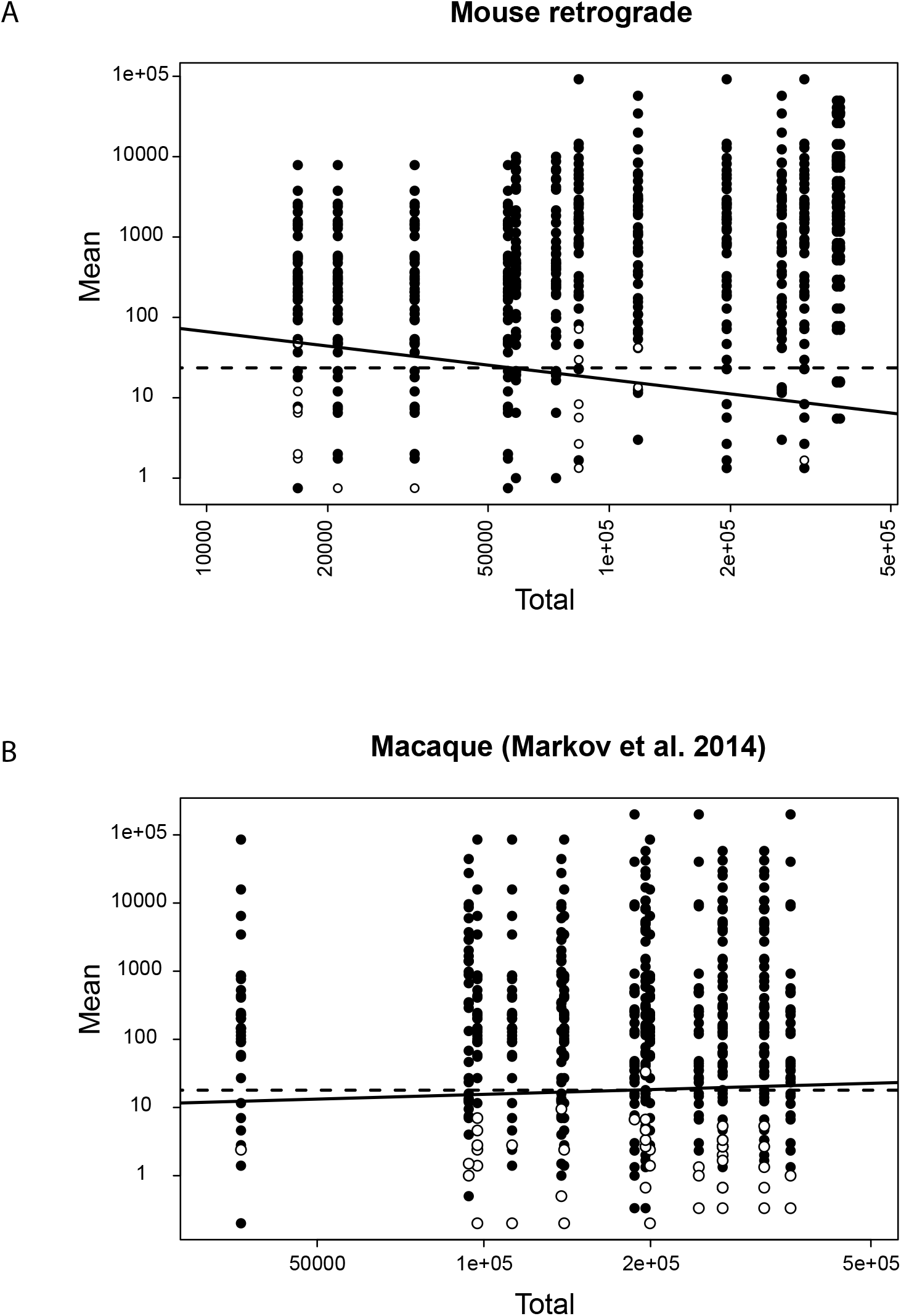
Consistency as a function of mean weight and size of injection for repeat injections across individuals. (**A**) Repeat injections in V1 (n=4); (**B**) LM (n=3). Black dots, projection present; white dots absent. Vertical axis, mean number of neurons per projection, horizontal axis injection size in terms of total number of neurons per injection. The solid lines correspond to a linear classifier from a logistic regression with the variables of both axes used as features for a probability of the presence of the projection at 95%. The dashed lines correspond to a similar criterion for which only the ordinate variable was used as a classification feature.

In Figure 4b, we perform a similar analysis on the macaque data reported by Markov et al (Markov et al., 2014) from repeat injections in areas V1, V2, V4 and 10. The solid and dashed lines correspond to the same models as in Figure 4a but fit to the macaque data. Again, the influence of the total size of the injection was not found to contribute significantly to the classifier. The dashed line corresponds to a mean of 18 neurons, slightly lower than the value found for the mouse data and close to the value of 10 estimated more informally in Markov et al. (Markov et al., 2014).

Verification of the model of data variability, estimation of dispersion and consistency as well as the generality of the lognormal distribution of weights and consistency justified the use of a single cortical injection of retrograde tracer in macaque to characterize the projection profile of an area (Markov et al., 2014). The present results show that this also holds for the retrograde labeling of cortical pathways in mouse.

### The mouse cortical connectome exhibits distinct connectivity profiles

Armed with a description of the distribution of the data, we tested whether there are signatures in the sets of projections to each area, as is the case in macaque (Markov et al., 2011). Alternatively, every individual might present its own sets of connections and weights. To decide this, examined each set of multiply injected areas to determine the minimum number of factors accounting for the systematic effects on the data.

For every group of repeat injections in V1 and LM (Figure 3 A, B), we modeled the number of labeled cells in the source areas as a function of two explanatory variables: the identity of areas and the identity of the injected individual. We fitted the data with generalized linear models (McCullagh and Nelder, 1989) and evaluated the influences of the explanatory variables over the models using Akaike’s Information Criterion (see Supplementary Information). The best resulting models were those for which individual differences appeared as unsystematic variability, i.e. without an interaction between the areas and the individual animals. The presence of such an interaction would have signified the presence of individual differences in connectivity profiles beyond the variability among animals. Its absence implies that quantitative connectivity profiles do not differ sufficiently across cases and therefore that a robust signature (connectivity profile) exists for each area.

To illustrate the relationship between the different levels of the statistical modeling, we performed a simulation of the expected experimental results under three different scenarios for the sampling distribution of the data: Poisson, Negative Binomial with *θ* = 7, and Geometric (i.e., Negative Binomial with *θ* = 1). In each case, we first simulated a log normal distribution of FLNe with mean and standard deviation based on one of our injections (red, green and blue curves in Figures 5A-C). Then we simulated 1000 repeats with dispersion specified according to each of the models. The results of these repeats are plotted as grey, semi-transparent points in Figures 5 A-C. Figure 5 D shows the SDs of the FLNe in the source areas plotted against the mean FLNe values, from the 1000 simulations, replicating the relation that we observed in Figure 3A, B.

**Figure 5.**
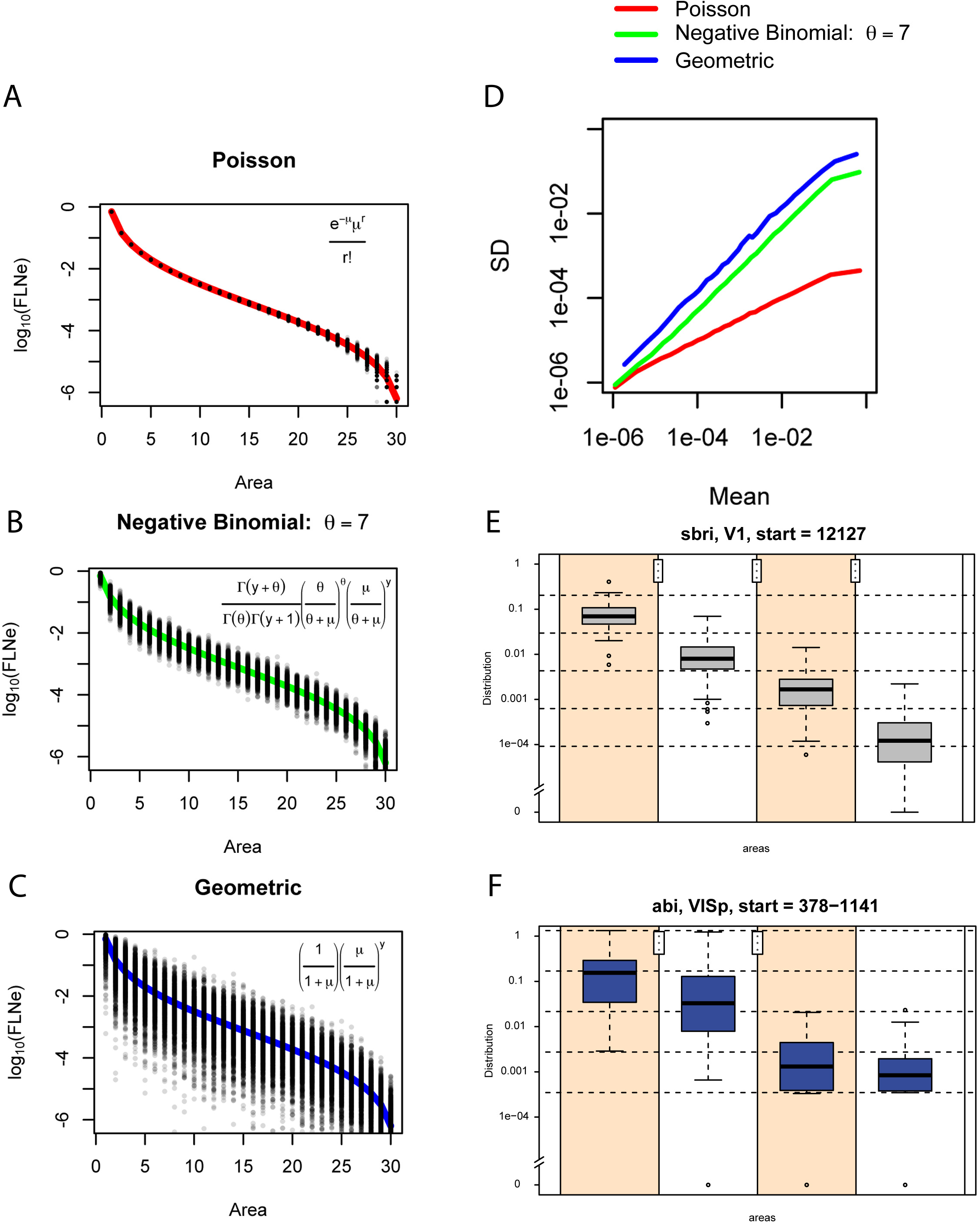
Relations of discrete probability distributions on log-normal FLNe distribution variabilities and connectivity profiles. In **A**, **B**, **C**, the hypothetical results of 1000 injections were simulated according to a Poisson, Negative Binomial (dispersion parameter theta = 7) and Geometric distributions (theta = 1). We considered 30 source areas with fixed mean FLNe spanning 6 orders of magnitude and evenly distributed along an average log-normal curve with the same mean and SD as one of our injections (red, green and blue, respectively, for the three distributions). Each mean FLNe was multiplied by 10^6^ to give the expected mean numbers of neurons in the source areas. These values were used for generating random counts from each of the distributions (Poisson, Negative Binomial and Geometric), as indicated in the insets of **A-C**. The set of random counts from every simulation were then normalized by their sum to transform them to simulated values of FLNe. This procedure was repeated 1000 times, thus revealing the expected spread of results from 1000 injections under each of the hypotheses. (**D**) The standard deviation is plotted as a function of the means calculated for the simulated injections from A, B and C with the colors indicating the distribution from which the calculations were made. (**E, F**) Example of the effect of overdispersion on the reliability of projections in present data and in Oh et al. 2014. In both plots a single injection in V1 (VISp, respectively) was taken and the areas were ordered according to their strengths. The difference between the log of the maximum and of the nonzero minimum was then divided into four intervals (delimited by dashed horizontal lines), and assigned the log of the FLNe to the corresponding intervals, forming four groups. Next, the strengths of the corresponding areas from the other repeats were used to obtain the boxplots. The stars represent the significance levels attained of the p values of one-sided permutation tests for each pair of consecutive groups, with the null hypothesis that the mean of the group on the left is larger than the mean of the group on the right. Notice that the present data are all restricted to the initial intervals (within the limits of the dashed horizontal lines), while the data from Oh et al. 2014 in all but one case cross these limits.

In the case of the Poisson distribution (Figure 5A), it is evident from the tight distribution of points about the ordered log normal curve, that the data would show a clear example of a connectivity profile or signature and an individual injection is likely to closely reflect the average behavior indicated by the red curve. At the other extreme is the geometric distribution in Figure 5C. Here, the high variability (2-3 orders of magnitude range of variation for each projection) largely obscures the systematic trend of the expected curve (blue). As shown in simulations by Scannell (Scannell et al., 2000), data distributed in this fashion would require an inordinate number of repeat injections to establish the average behavior of the curve with sufficient precision. Note that individual injections could follow any arbitrary path through the point cloud, so their value in establishing an areal profile would be nearly meaningless. Statistical analysis of a small number of injections would likely lead to the conclusion of individual differences in the profile for a single injection site, that is the presence of a statistically significant interaction between area and brain injected. Figure 5B displays the simulated results from a Negative Binomial distribution, which with a dispersion parameter similar to that found in retrograde labeling in macaque and mouse, falls in between the two other distributions. However, with the variation of individual injections being only 1 order of magnitude, far less than the span of the ordered log normal curve, single injections are much more representative of the average curve than for geometrically distributed data. As shown in our data, the variation among animals is not sufficient to reject the proposition that projection profiles from different animals are the same.

In order to illustrate the effects of a geometric distribution (Figure 5C) of the anterograde data and negative binomial distributions (Figure 5B) for the retrograde data on connectivity profiles, we show the box plot analysis for area V1 for the present data (Figure 5E) and for the Oh et al., 2014 data (Figure 5F). This shows that the negative binomial distribution corresponds to a significantly more demarcated connectivity profile compared to the data with a geometric distribution in figure 5F from Oh et al., 2014.

### The ultra-high density of the cortical graph

The number of connections in the full interareal network (FIN), expressed as a fraction of the number of possible connections, defines the graph density which is a fundamental measure of the graph’s overall connectedness, extensively used in network science and also in earlier analyses of cortical connectivity (Markov et al., 2013b; Markov et al., 2014; Sporns and Zwi, 2004). Referring to the weighted connectivity matrix in Figure 6, but employing the corresponding binary connectivity matrix, below we infer the density of the FIN (Janson et al., 2000; Markov et al., 2014; Newman, 2010); for the full FLN weights data see Supplementary Table 1. Based on an atlas of 47 areas (41 areas of which SSp was divided into 7 subareas, Figure S1B), the FIN contains *N_FIN_* = 47 cortical areas that represent the nodes of the *G_47x47_* graph. The directed edges of the FIN correspond to directed connections between nodes, based on the fraction of labeled neurons. Our analysis of the FIN makes use of the *G_19x47_* directed subgraph of projections within FIN, which reveals all the in-degrees of the injected 19 nodes. It also makes use of the *G_19x19_ edge-complete* subgraph of FIN, corresponding to the connections among just the 19 injected areas. Both *G_19x47_* and *G_19x19_* contain complete information about the status of their edges and would not be influenced by injections into additional areas elsewhere in the cortex. Given that the 19 injected areas are widely distributed across the cortex (see Fig. S1B), the *G_19x19_* subgraph is likely to reflect major characteristics of the FIN.

**Figure 6.**
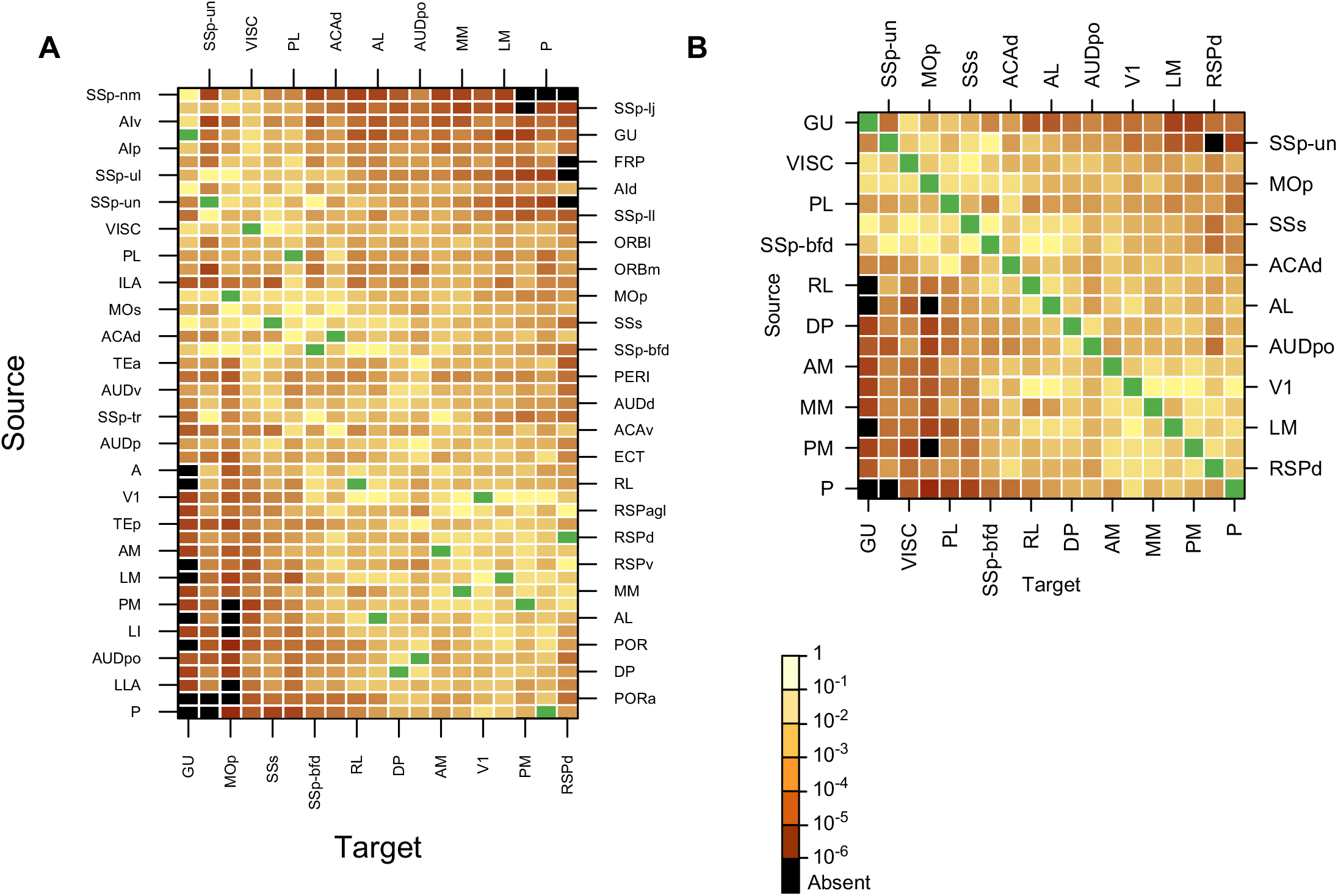
Weighted connectivity matrix. The color shows the strength of the projection as indicated by the color bar with black corresponding to absent connections and green for the intrinsic projections that are not included here. (**A**) Each row represents one of the 47 source areas; each column represents one of the 19 injected target areas. Note that the SSp-bfd and SSp-un subfields are listed as separate areas. The row and column ordering was determined by a clustering algorithm based on similarity of the input and output profiles between areas (see Methods). (**B**) A weighted connectivity matrix for the 19 x 19 subgraph. For multiple injections, shading is based on arithmetic mean values of FLN.

The density of a directed graph is given by the ratio *ρ*=*M*/[*N*(*N*-1)] between the number of directed edges (links) *M* of the graph and the maximum possible number of directed links, *N*(*N*-1), where *N* is the number of nodes in the graph. The *G_19x19_* graph has *M*= 334 (binary) directed links from the maximum possible of *N*(*N*-1) = 342, and therefore it is strongly interconnected, with a very high graph density of *ρ*=0.97 (97%). Because it is an edge complete subgraph of FIN, the density of *G_19x19_* is expected to be comparable to that of the FIN.

The in-degrees of the *G_19x47_* graph (i.e., the number of source areas projecting to each of 19 target areas), range from 38 to 46 with a mean of <k>_in_ = 44.8 (Figure S4B); their distribution is asymmetrical and NOT concentrated around the mean, but instead strikingly close to the maximum (Figure S4B, top right). The density of the FIN can be estimated as follows. Every directed edge is an in-link to some node so that the total number of edges *M_FIN_* equals the total number of in-links in the FIN. We lack data on the in-links to nodes that were not injected, but we can assume that they are characterized by the same average in-degree as for the 19 injected nodes. Assuming M_FIN_ ~ <k>in N_FIN_ = 2107 for the FIN (*G_47x47_*) leads to the prediction that ρ_FIN_ = M_FIN_ / [N_FIN_ (N_FIN_ − 1)] ~ <k>in / (N_FIN_ − 1) ~ 0.97).

A dominating set analysis on *G_19x47_* provides further evidence that the FIN is indeed dense (Figure S4A). In graph theory, a subset *D* of nodes of a graph *G* with node set *V* is said to be dominating G, if all elements of *V* have a link to at least one node in *D* (Kulli and Sigarkanti, 1991). Here we modify this definition slightly by saying that *D dominates x% of the nodes of G*, if an *x*% of *all* nodes in *V* are linked to one or more nodes in *D*. The *x*%=100% corresponds to “full” domination. This definition includes also nodes from *D*. The Minimum Dominating Set (MDS) *Dmin* is defined as the one that fully dominates *G* and it has the smallest size (number of nodes). For all sets of 2 target area combinations from the 19 target parcels (171 pairs), 92% of them dominate 100% of the 47 parcels (Table S1). Thirteen parcels out of 19 (68%) are fully connected.

A low MDS indicates either a very dense graph, or a scale-free graph (dominated by its hubs) (Barabasi and Albert, 1999). Both the in-degree distribution (Supplementary Figure 4B, top right) *and* the fact that slightly increasing the size of dominating sets to include 3, 4 and more nodes quickly increases their number, shows incompatibility with a scale-free graph. The high density by itself is *incompatible* with a scale-free network as shown in (Del Genio et al., 2011). For area triples, there are 965 dominating sets (99.6% of 969). Moreover, all combinations of 4 sites (out of 19, i.e., 3876) will dominate 100% of all the areas (Table S1). Additional injections can add links but not new nodes, and therefore can only enhance these strong domination effects.

These results show that retrograde tracers using a deterministic experimental design reveal a dense cortical matrix which suggests an ultradense cortical network of 97%, considerably higher than the densities found using a computational model based on anterograde tracers (Oh et al., 2014). Comparison of the present results with this work provides some measure of the differences in the empirical results in both studies and suggests that modeling assumptions for determining the location of injection might be partly responsible for the differences in the reported densities (see Discussion section).

## Discussion

PVtdT-expressing mice allowed accurate areal parcellation in individual animals used for tracer injections, as reflected in the spatial map of labeled projection neurons shown in Figure 2. The exploitation of quantitative retrograde tracing in these flat maps was an important part of our experimental design aimed at minimizing inter-animal sampling errors. Experimental design of the present study made it possible to verify that the well-defined uptake zone of the injection site (Markov et al., 2014) was entirely confined to the intended area. Modeling of FLNe variance in repeat injections across animals allowed exclusion of a geometric distribution in favor of a negative binomial distribution (Figure 3). Ranking FLNe values revealed lognormal distributions spanning five orders of magnitude with estimated 95% confidence intervals that satisfactorily contained the mean values (Figure 3). In addition, the consistency analysis of repeat injections showed that means of excess of twenty-four neurons were consistent across injections (Figure 4). Finally, the density of the matrix of the cortical subgraph studied here suggests that the full matrix may have a density of 97% (Figure 5).

This density is considerably higher than the maximum 53% reported in the probabilistic mouse connectivity matrix (Oh et al., 2014). High connection density may be in part due to spillage of DY beyond the borders of areas or grey matter, and/or uptake by fibers of passage and damaged axons (Keizer et al., 1983). None of these caveats can be ruled out with certainty. However, based on images like those in Figure S2, which reveal the extent and location of the injection sites, and eliminating the cases which showed obvious involvement of more than one area, we are confident that undetected spillage across areal borders contributed minimally to the high connection density. Similarly, cases with breaches of the unmistakable grey/white matter border were eliminated from the analysis and therefore this is unlikely to be a contributing factor. On the other hand, labeling of area-to area-projecting neurons through spurious uptake of DY by deep layer axons is a potential concern. However, based on the tight and topographically precise distribution of DY labeled cells (Figure 2, Figure S2B, C,) the impact seems minor. The differential labeling of dorsal and ventral stream areas after injections of V1, which may label damaged area-to area axons in transit further supports this conclusion.

In the study by Oh et al. 2014, the location of injection sites were inferred from a template, and the published database indicates that 76% of injections in isocortex involved multiple cortical areas (at least 2 and up to 9 areas). In that study, connections resulting from mixed injections were disentangled and the strength of connections between individual areas estimated algorithmically. The low density estimation of the cortical graph density in the Oh et al., 2014 study could be due to the computational procedure used (Ypma and Bullmore, 2016), as we show in the next section.

### Comparison of weight distribution in the present results with previous studies

While many of the anterograde tracer injections in isocortex in the Oh et al., 2014 study were mixed injections, 26 of them were restricted to single areas (14 areas injected in total). The authors derived from the mixed and non-mixed injections the full matrix of interareal connections, which we will name computational data. In contrast, the connections from injections confined to single areas (non-mixed injections) will be named raw data. Comparison of raw connection strengths with those obtained from the computational model allows appraisal of the modeling assumptions used in that study. Figure 7A. shows the agreement between the raw and the computed connections, with only 65% of them being true positives or true negatives, while 33% are false negatives (found in raw but not in computed) and 2% false positives (found in computed but not in raw). Moreover, the squared correlation between the true positives is modest, of only 0.58. Examination in Figure 7B of the weight distribution of the raw and computed data gives further insight into how the computational algorithm of the Oh et al., 2014 transforms the raw data. Here the blue bars are the connections that result from the raw non-mixed injections. The red bars are the computed set of connections corresponding to the raw connections. The white bars are the connections that are present in the raw data but which are missing in the computed data. The computed data equivalent to the 14 injected areas returns 314 connections, significantly less than the 478 connections observed in the raw data. The 164 connections that are present in the raw data but absent in the computed (white bars), while predominantly weak are nevertheless found throughout the full range of weights. This figure using a log scale for connection strength suggests that the Oh et al., 2014 raw data does not follow the same lognormal distribution as the computed data.

**Figure 7.**
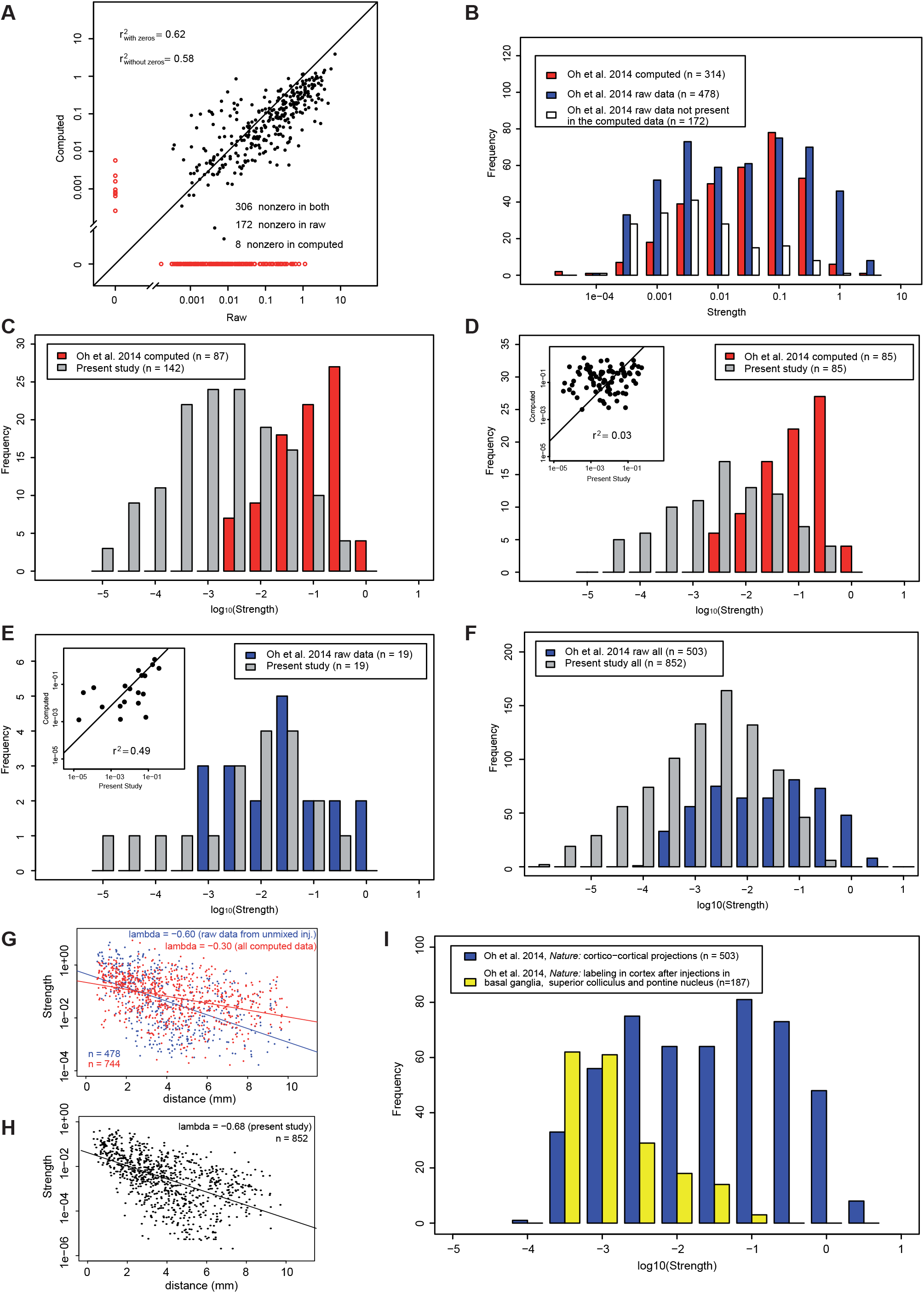
The data in the present study shows some similarity to the raw data in Oh et al. 2014, but not to the computed data. (**A**) Correlation between the raw and computed data in Oh et al., 2014, zero values shown in red. (**B**) Distributions of the raw data and the corresponding connections in the computed data for the 14 areas which received unmixed injections in Oh et al., 2014; red bars, computed data, grey bars, raw data, white bars, nonzero connections in the raw data, but zero in the computed. (**C**) Distribution of strengths of connections for areas which are homologous in Oh et al. 2014, computed data (red) and present study (grey). Source areas: ACAd, ACAv, AId, AIp, AIv, ECT, GU, ILA, MOp, MOs, ORBl, ORBm, ORBvl, PERI, PL, RSPd, RSPv, SSp-bfd, V1; target areas, ACAd, GU, ILA, MOp, MOs, RSPd, SSp-bfd, V1. (**D**) Same as in C, but considering only projections which are nonzero in both sets. Insert, correlation diagram. (**E**) Distribution of strengths of connections for areas which are homologous and non-zero both in Oh et al. 2014, raw data (blue, the 14 areas which received unmixed injections) and present study (grey). Insert, correlation diagram. Source areas: MOp, SSp-bfd, V1; target areas: ACAd, GU, ILA, MOp, MOs, RSPd, SSp-bfd, V1. (**F**) Distribution of connection strengths for the full data set in present study (gray bars) compared to raw data in Oh et al., 2014 shown in panel B. (**G**) Distribution of projection lengths in Oh et al., 2014, raw data (blue) and computed data (red). (**H**) Distribution of projection lengths in present study. Notice that the spatial constant is close to the one in raw data in G. (**I**) Comparison of cortical labeling in Oh et al., 2014 follow anterograde tracer injections in the superior colliculus, pontine nucleus and basal ganglia with label obtained following cortical injections.

In Figure S3, nineteen of the 41 cortical areas in an adapted atlas from the present study correspond to areas in the atlas used by Oh et al., 2014, allowing some direct comparison between the two studies (listed in Fig 7 legend). The present study found 142 connections, which contrasts with the 87 connections computed for these areas by Oh et al., 2014 using anterograde tracers. Figure 7C shows the set of connections shared by the two studies (same source area and same target area). We can observe that the computed data from the Oh et al., 2014 exhibits a lognormal distribution tightly restricted to the top three orders of magnitude compared to the broad lognormal distribution in the present study. Figure 7D provides a more direct comparison, by contrasting only those connections that are non-zero in both studies. They differ significantly in their weight range, and more importantly show no correlation (insert Figure 7D). These findings confirm that the algorithm used by Oh et al., 2014 to disentangle connections is leading to significant transformations by reducing the number of connections and influencing their distribution. This is further supported by comparing the 19 common connections found in the raw data in Oh et al., 2014 with their homologues in the present data (Figure 8E). Although limited this suggests that compared to the computed data, the raw data in the Oh et al., 2014 shows an improved overlap in the weights and has a modest correlation with data from the present study (see Figure 8E insert). However, Figure 8F shows that the raw data of the Oh et al., 2014 possesses a narrow range of weights compared to the present study.

**Figure 8.**
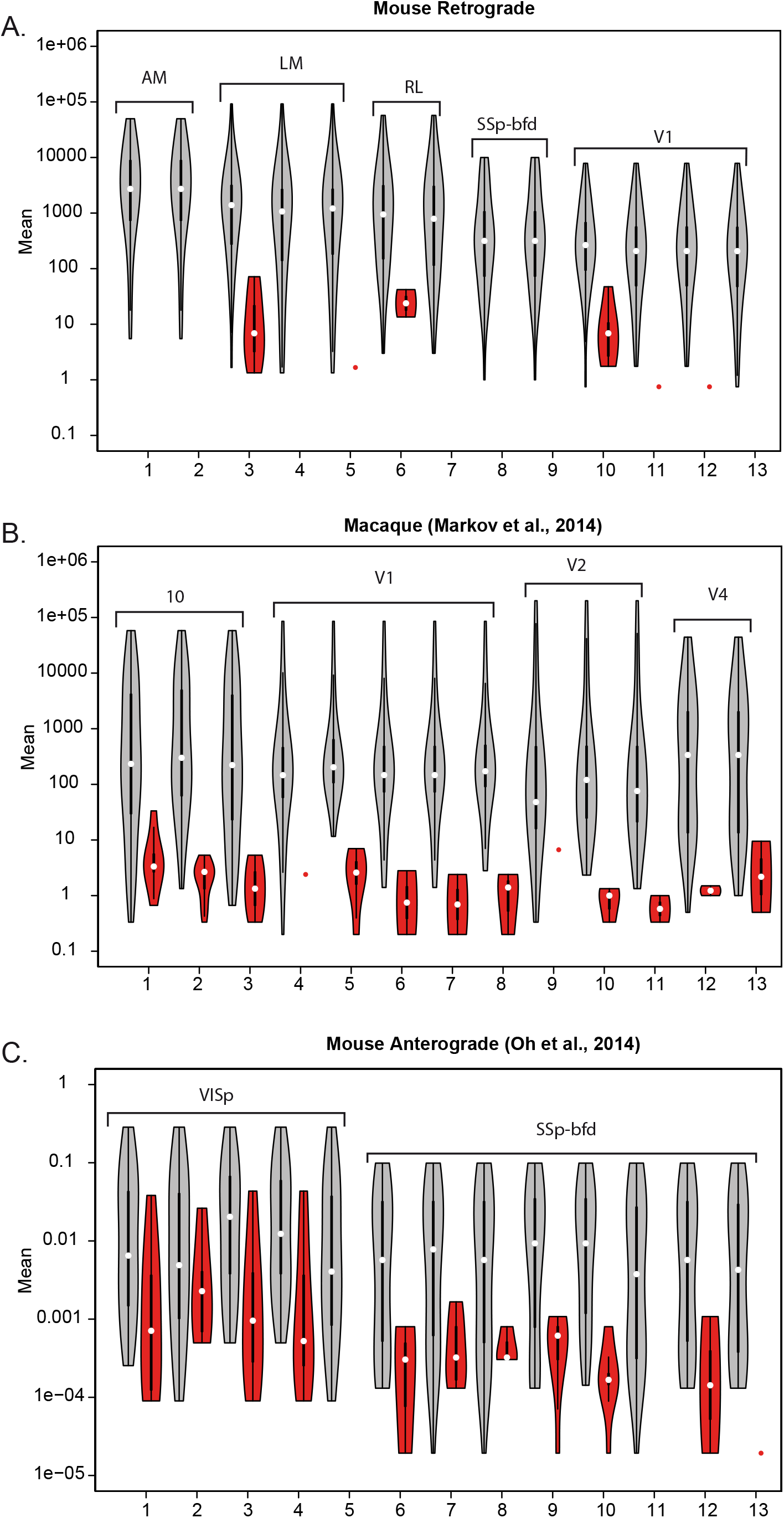
High consistency of retrograde compared to anterograde data. Violin plot of means of projections consistent across repetitions (gray), of inconsistent projections (red). (**A**) Mouse retrograde tracing data repeat injections in areas AM, LM, RL, SSp-bfd and V1; (**B**) Macaque retrograde tracing data from repeat injections in areas V1, V2, V4 and 10 (Markov et al., 2014). (**C**) Mouse raw anterograde data where repeat injections were restricted to single areas (VISp, SSp-bfd) (Oh et al., 2014).

Elsewhere we have shown that the weight-distance relationship is a cardinal feature of the connectome that in both mouse and monkey is related to geometrical features such as the motif distribution of the cortical graph (Ercsey-Ravasz et al., 2013; Horvat et al., 2016). In Figure 7G we compare the decline of weight with distance in the computed and raw cortico-cortical connections from Oh et al., 2014. This shows only a very modest slope for the computed data, but a slope in the raw data comparable to that found in the present study (Figure 7H). The slope of -0.68 obtained in the present study is similar to that obtained in Horvat et al., (2016). Finally, in the Oh et al., 2014 study, the raw data from the unmixed injection return a density of 93% which is consistent with our observed density of 97%.

In anterograde tracing it is challenging to distinguish between pre-terminal axons and boutons and to exclude labeled fibers of passage from the analysis. Further, there is an important difference in scale explored by the two tracers; anterograde is subcellular as it reveals individual boutons with as many as tens of thousands per axon. In contrast, retrograde tracing is at the single cell level. These differences are compounded by the difficulty in quantification of the anterograde signal using optical density and by the notorious problem of anterograde tracers, which occasionally label neurons retrogradely, and with them their collaterals both in the injected area and other areas to which the collaterals project (Reiner et al., 2000). Estimates of terminal densities from anterograde tracers using optical density measurements where axons of passage can introduce a significant bias may be less accurate than counts of labeled neurons following retrograde tracers. Since the observations of LeVay and Sherk (LeVay and Sherk, 1983), anterograde tracers have been shown to lead to retrograde labeling, and this includes the viral tracers used in the Oh et al., 2014 study (Wang et al., 2014). This can lead to labeling of local collaterals of the retrogradely labeled cells and could explain the secondary spurious anterograde labeling observed in the Oh et al., 2014 study, where anterograde injections in the pontine nucleus and superior colliculus led to levels of labeling in cortex comparable to those reported for cortico-cortical projections (see Figure 7I, and Table S2).

### Comparison of variability of deterministic connectivity maps in mouse and monkey

Variation in connectivity in addition to impacting on the connectivity profile will also influence the consistency of connections in a particular pathway i.e. the extent to which connections are found to be present across animals. In Figure 8 are shown violin plots for connections that are found consistently across injections (gray) and those that are absent from a given injection (red). In the present study there is a high degree of consistency across injections, inconsistent connections were only found in significant numbers in 3 of the 13 repeats and further were restricted to weak projections (Figure 8A). The restriction of inconsistent connections to weak connections was also found in macaque. The macaque data in Markov et al. (Markov et al., 2014) differed from the mouse in that significant levels of inconsistency were found in ten of the 13 repeat injections (Figure 8B). The higher incidence of inconsistency in macaque could be related to the partial sampling in the macaque study. The high levels of consistency in the retrograde tracer studies of mouse and macaque differ from that observed in the Oh et al., 2014 anterograde data where inconsistent projections were found across nearly all injections and at much larger weight values (Figure 8C).

Various considerations could suggest species differences in the variability of cortical connectivity. The number of cortical areas in an ancestral mammal is not known, but evolution evidently led to an increase in the number of cortical areas in the primate lineage (especially humans) compared to the rodent/mouse lineage (Striedter, 2005). Evolutionary changes include a relative increase in the extent of the cortical mantle that is referred to as association cortex in primates (i.e. cortex outside of the early sensory/motor areas). Cortical development is known to be under both intrinsic and extrinsic factors. Environmental factors that are known to influence the development of the cortex (Kennedy and Dehay, 1993; O'Leary et al., 2007) could potentially have a differential impact on arealization and variability in cortical connectivity in mouse and macaque (Buckner and Krienen, 2013). Further, the laboratory mouse is a highly inbred strain that could exhibit less phenotypic variability than the macaque.

Retrograde tracers are used to reveal the set of source areas that project to a given target area. Together, the specific set and the strength or weights of each projection determines the connectivity profile of a given area which, coupled with the intrinsic connectivity of the target area, are presumably critical for its functional characteristics (Bressler and Menon, 2010; Markov et al., 2013a; Passingham et al., 2002).

Many of the connections linking cortical areas show very low weights which lead us to speculate that weak connections in macaque might nonetheless play an important role in shaping cortical function (Markov et al., 2013a; Markov et al., 2013b; Markov et al., 2011). These observations are in accordance with the weak link concept of social and ageing biological networks in which the loss of sparse connections render systems unstable (Csermely, 2006; Granovetter, 1973) as well as the theoretical analysis of the macaque cortical connectome (Goulas et al., 2015) and recent imaging data in human (Bassett and Bullmore, 2016). How consistent are these weak connections? In macaque almost all connections greater than a mean of ten neurons were consistent (Markov et al., 2014). In the current study, our more rigorous statistical quantification of this indicated a 95% connection probability for a mean of 18 or more neurons. The present retrograde tracing results in mouse show marginally higher inconsistency compared to monkey, but nevertheless considerably more consistent than the anterograde study in the Oh et al., 2014 where reported inconsistency was more frequent (Figure 8). Hence, the present study in the mouse confirms findings in macaque and argue that weak connections are consistent and need to be considered in large-scale models of the brain.

### Density of the mouse cortical graph and its implications

Graph density is an important measure of the level of connectivity in a network. While most empirical networks (such as social networks, technological/IT networks, infrastructure networks, gene regulatory networks, metabolic networks, protein interaction networks, etc.) are sparse, cortical interareal networks, surprisingly, form high density graphs. A graph is referred to as sparse when the number of links is of the same order as the number of nodes: for example, for a network of 19 nodes that would imply on the order of 20 ~ 50 directed links, not the 334 found in the present study. The high-density character of the cortical network first observed in the macaque, resulted in a graph with a density of 66% (Markov et al., 2011). The current data, obtained with the same deterministic approach shows that the mouse interareal network is ultradense with a density of 97%, corresponding to a nearly complete graph. This implies that almost all area pairs have direct connectivity, *both ways*, suggesting high integration of information across the whole cortical hierarchy. While the binary picture eliminates almost all specificity at such high densities, once we take into account the weights of the connections, a very different picture emerges. Even at the most basic level, inspection of the weighted connectivity matrix in Figure 6B demonstrates striking asymmetries in many bidirectional connectivity strengths, showing that the *G_19x19_* is a directed graph with strong weight specificity. This is also evident from comparison of individual tracer injections, for example, V1 injections show sparse labeling of somatosensory sub-areas, whereas somatosensory injections show much stronger labeling in V1. This means that in order to understand processing and information flow in such networks one must use methods that exploit the weighted nature of connections (Barrat et al., 2004; Newman, 2004), and the geometrical and morphological features of the areas within the cortical plate.

The mouse brain measures less than 8 mm in length but despite its small size it has become an increasingly important model for investigation of higher functions of the cortex. The sophistication of current investigations of the neuronal basis of higher function in mouse has allowed unprecedented progress in neuroscience. The fact that the mouse cortex has a graph density of 97% has an undeniable impact on the way we understand the relationship between the structure and function of the cortex. The mouse cortical graph can achieve a high functional specificity despite its high density because it has pronounced connectivity profiles. To understand the consequences of the high density network of the mouse brain relative to the sparser networks of larger brains will require developing weighted network comparisons, which are not a well-established area in current network science.

## Acknowledgements

This work was supported LABEX CORTEX (ANR-11-LABX-0042) of Université de Lyon (ANR-11-IDEX-0007) operated by the French National Research Agency (ANR) (H.K.), ANR-11-BSV4-501, CORE-NETS (H.K.), ANR-14-CE13-0033, ARCHI-CORE (H.K.), ANR-15-CE32-0016, CORNET (H.K.), Fédération des Aveugles de France (R.G.), McDonnell Center for Systems Neuroscience (A.B. and R.G.) and NIH grant RO1 EY022090 (A.B).

## References

Barabasi, A.L., and Albert, R. (1999). Emergence of scaling in random networks. Science 286, 509–512.

Barrat, A., Barthelemy, M., Pastor-Satorras, R., and Vespignani, A. (2004). The architecture of complex weighted networks. Proc Natl Acad Sci U S A 101, 3747–3752.

Bassett, D.S., and Bullmore, E.T. (2016). Small-World Brain Networks Revisited. Neuroscientist.

Bressler, S.L., and Menon, V. (2010). Large-scale brain networks in cognition: emerging methods and principles. Trends Cogn Sci 14, 277–290.

Buckner, R.L., and Krienen, F.M. (2013). The evolution of distributed association networks in the human brain. Trends Cogn Sci 17, 648–665.

Bullmore, E., and Sporns, O. (2012). The economy of brain network organization. Nat Rev Neurosci 13, 336–349.

Buzsaki, G., and Mizuseki, K. (2014). The log-dynamic brain: how skewed distributions affect network operations. Nat Rev Neurosci 15, 264–278.

Carandini, M., and Churchland, A.K. (2013). Probing perceptual decisions in rodents. Nat Neurosci 16, 824–831.

Csermely, P. (2006). Weak Links: Stabilizers of complex systems from protein to social networks (Berlin: Springer).

Del Genio, C.I., Gross, T., and Bassler, K.E. (2011). All scale-free networks are sparse. Phys Rev Lett 107, 178701.

Dong, H.W. (2008). The Allen reference atlas: A digital color brain atlas of the C57B1/6J male mouse (John Wiley & Sons Inc).

Ercsey-Ravasz, M., Markov, N.T., Lamy, C., Van Essen, D.C., Knoblauch, K., Toroczkai, Z., and Kennedy, H. (2013). A predictive network model of cerebral cortical connectivity based on a distance rule. Neuron 80, 184–197.

Felleman, D.J., and Van Essen, D.C. (1991). Distributed hierarchical processing in the primate cerebral cortex. Cereb Cortex 1, 1–47.

Ferezou, I., Haiss, F., Gentet, L.J., Aronoff, R., Weber, B., and Petersen, C.C. (2007). Spatiotemporal dynamics of cortical sensorimotor integration in behaving mice. Neuron 56, 907–923.

Garrett, M.E., Nauhaus, I., Marshel, J.H., and Callaway, E.M. (2014). Topography and areal organization of mouse visual cortex. J Neurosci 34, 12587–12600.

Glasser, M.F., Smith, S.M., Marcus, D.S., Andersson, J.L., Auerbach, E.J., Behrens, T.E., Coalson, T.S., Harms, M.P., Jenkinson, M., Moeller, S., Robinson, E.C., Sotiropoulos, S.N., Xu, J., Yacoub, E., Ugurbil, K., and Van Essen, D.C. (2016). The Human Connectome Project's neuroimaging approach. Nat Neurosci 19, 1175–1187.

Goulas, A., Schaefer, A., and Margulies, D.S. (2015). The strength of weak connections in the macaque cortico-cortical network. Brain Struct Funct 220, 2939–2951.

Granovetter, M.S. (1973). The strength of weak ties. Am J Sociology 78, 1360–1380.

Hilbe, J.M. (2007). Negative Binomial Regression (Cambridge: Cambridge University Press).

Hippenmeyer, S., Vrieseling, E., Sigrist, M., Portmann, T., Laengle, C., Ladle, D.R., and Arber, S. (2005). A developmental switch in the response of DRG neurons to ETS transcription factor signaling. PLoS Biol 3, e159.

Horvat, S., Gamanut, R., Ercsey-Ravasz, M., Magrou, L., Gamanut, B., Van Essen, D.C., Burkhalter, A., Knoblauch, K., Toroczkai, Z., and Kennedy, H. (2016). Spatial Embedding and Wiring Cost Constrain the Functional Layout of the Cortical Network of Rodents and Primates. PLoS Biol 14, e1002512.

Janson, S., Luczak, T., and Rucinski, A. (2000). Random graphs (Wiley-Interscience).

Keizer, K., Kuypers, H.G.J.M., Huisman, A.M., and Dann, O. (1983). Diamidino Yellow dihydrochloride (DY 2HCl): a new fluorescent retrograde neuronal tracer, which migrates only very slowly out of the cell. Exp Brain Res 51, 179–191.

Kennedy, H., and Dehay, C. (1993). Cortical specification of mice and men. Cereb Cortex 3, 27–35.

Kennedy, H., Knoblauch, K., and Toroczkai, Z. (2013). Why data coherence and quality is critical for understanding interareal cortical networks. Neuroimage 80, 37–45.

Kim, H., Ahrlund-Richter, S., Wang, X., Deisseroth, K., and Carlen, M. (2016). Prefrontal Parvalbumin Neurons in Control of Attention. Cell 164, 208–218.

Krubitzer, L.A., and Seelke, A.M. (2012). Cortical evolution in mammals: the bane and beauty of phenotypic variability. Proc Natl Acad Sci U S A 109 Suppl 1, 10647–10654.

Kulli, V.R., and Sigarkanti, S.C. (1991). Inverse domination in graphs. Nat Acad Sci Letters 14, 473–475.

LeVay, S., and Sherk, H. (1983). Retrograde transport of [3H]proline: a widespread phenomenon in the central nervous system. Brain Res 271, 131–134.

Li, N., Daie, K., Svoboda, K., and Druckmann, S. (2016). Robust neuronal dynamics in premotor cortex during motor planning. Nature 532, 459–464.

Lindsey, J.K. (1999). Models Repeated Measurements, 2nd edn (Oxford Oxford University Press).

MacNeil, M.A., Einstein, G., and Payne, B.R. (1997). Transgeniculate signal transmission to middle suprasylvian cortex in intact cats and following early removal of areas 17 and 18: a morphological study. Exp Brain Res 114, 11–23.

Madisen, L., Zwingman, T.A., Sunkin, S.M., Oh, S.W., Zariwala, H.A., Gu, H., Ng, L.L., Palmiter, R.D., Hawrylycz, M.J., Jones, A.R., Lein, E.S., and Zeng, H. (2010). A robust and high-throughput Cre reporting and characterization system for the whole mouse brain. Nat Neurosci 13, 133–140.

Manita, S., Suzuki, T., Homma, C., Matsumoto, T., Odagawa, M., Yamada, K., Ota, K., Matsubara, C., Inutsuka, A., Sato, M., Ohkura, M., Yamanaka, A., Yanagawa, Y., Nakai, J., Hayashi, Y., Larkum, M.E., and Murayama, M. (2015). A Top-Down Cortical Circuit for Accurate Sensory Perception. Neuron 86, 1304–1316.

Markov, N.T., Ercsey-Ravasz, M., Lamy, C., Ribeiro Gomes, A.R., Magrou, L., Misery, P., Giroud, P., Barone, P., Dehay, C., Toroczkai, Z., Knoblauch, K., Van Essen, D.C., and Kennedy, H. (2013a). The role of long-range connections on the specificity of the macaque interareal cortical network. Proc Natl Acad Sci U S A 110, 5187–5192.

Markov, N.T., Ercsey-Ravasz, M., Van Essen, D.C., Knoblauch, K., Toroczkai, Z., and Kennedy, H. (2013b). Cortical high-density counter-stream architectures. Science 342, 1238406.

Markov, N.T., Ercsey-Ravasz, M.M., Ribeiro Gomes, A.R., Lamy, C., Magrou, L., Vezoli, J., Misery, P., Falchier, A., Quilodran, R., Gariel, M.A., Sallet, J., Gamanut, R., Huissoud, C., Clavagnier, S., Giroud, P., Sappey-Marinier, D., Barone, P., Dehay, C., Toroczkai, Z., Knoblauch, K., Van Essen, D.C., and Kennedy, H. (2014). A weighted and directed interareal connectivity matrix for macaque cerebral cortex. Cereb Cortex 24, 17–36.

Markov, N.T., Misery, P., Falchier, A., Lamy, C., Vezoli, J., Quilodran, R., Gariel, M.A., Giroud, P., Ercsey-Ravasz, M., Pilaz, L.J., Huissoud, C., Barone, P., Dehay, C., Toroczkai, Z., Van Essen, D.C., Kennedy, H., and Knoblauch, K. (2011). Weight Consistency Specifies Regularities of Macaque Cortical Networks. Cereb Cortex 21, 1254–1272.

Marshel, J.H., Garrett, M.E., Nauhaus, I., and Callaway, E.M. (2011). Functional specialization of seven mouse visual cortical areas. Neuron 72, 1040–1054.

McCullagh, P., and Nelder, J.A. (1989). Generalized linear models, 2nd edn (Boca Raton: Chapman & Hall/CRC).

Mease, R.A., Metz, M., and Groh, A. (2016). Cortical Sensory Responses Are Enhanced by the Higher-Order Thalamus. Cell Rep 14, 208–215.

Musil, S.Y., and Olson, C.R. (1988a). Organization of cortical and subcortical projections to anterior cingulate cortex in the cat. J Comp Neurol 272, 203–218.

Musil, S.Y., and Olson, C.R. (1988b). Organization of cortical and subcortical projections to medial prefrontal cortex in the cat. J Comp Neurol 272, 219–241.

Newman, M.E.J. (2004). Analysis of weighted networks. PhysRev E 70, 056131.

Newman, M.E.J. (2010). Networks: an introduction (Oxford University Press).

O'Leary, D.D., Chou, S.J., and Sahara, S. (2007). Area patterning of the mammalian cortex. Neuron 56, 252–269.

Oh, S.W., Harris, J.A., Ng, L., Winslow, B., Cain, N., Mihalas, S., Wang, Q., Lau, C., Kuan, L., Henry, A.M., Mortrud, M.T., Ouellette, B., Nguyen, T.N., Sorensen, S.A., Slaughterbeck, C.R., Wakeman, W., Li, Y., Feng, D., Ho, A., Nicholas, E., Hirokawa, K.E., Bohn, P., Joines, K.M., Peng, H., Hawrylycz, M.J., Phillips, J.W., Hohmann, J.G., Wohnoutka, P., Gerfen, C.R., Koch, C., Bernard, A., Dang, C., Jones, A.R., and Zeng, H. (2014). A mesoscale connectome of the mouse brain. Nature 508, 207–214.

Olson, C.R., and Musil, S.Y. (1992). Topographic organization of cortical and subcortical projections to posterior cingulate cortex in the cat: evidence for somatic, ocular, and complex subregions. J Comp Neurol 324, 237–260.

Passingham, R.E., Stephan, K.E., and Kotter, R. (2002). The anatomical basis of functional localization in the cortex. Nat Rev Neurosci 3, 606–616.

Qi, H.X., and Kaas, J.H. (2004). Myelin stains reveal an anatomical framework for the representation of the digits in somatosensory area 3b of macaque monkeys. J Comp Neurol 477, 172–187.

Reiner, A., Veenman, C.L., Medina, L., Jiao, Y., Del Mar, N., and Honig, M.G. (2000). Pathway tracing using biotinylated dextran amines. J Neurosci Methods 103, 23–37.

Ringo, J.L. (1991). Neuronal interconnection as a function of brain size. Brain Behav Evol 38, 1–6.

Saleem, K.S., Logothetis, N.K. (2012). Atlas of the rhesus monkey brain (Amsterdam: Elsevier, Academic Press).

Scannell, J.W., Grant, S., Payne, B.R., and Baddeley, R. (2000). On variability in the density of corticocortical and thalamocortical connections. Philos Trans R Soc Lond B Biol Sci 355, 21–35.

Sherman, S.M. (2016). Thalamus plays a central role in ongoing cortical functioning. Nat Neurosci 19, 533–541.

Sincich, L.C., Adams, D.L., and Horton, J.C. (2003). Complete flatmounting of the macaque cerebral cortex. Vis Neurosci 20, 663–686.

Sporns, O., and Zwi, J.D. (2004). The small world of the cerebral cortex. Neuroinformatics 2, 145–162.

Striedter, G.F. (2005). Principles of brain evolution (Sunderland, MA: Sinauer Associates).

Van Essen, D.C., Glasser, M.F., Dierker, D., Harwell, J., and Coalson, T. (2012). Parcellations and hemispheric asymmetries of human cerebral cortex analyzed on surface-based atalses. Cereb Cortex 22, 2241–2262.

Venables, W.N., and Ripley, B.D. (2002). Modern Applied Statistics with S, 4th edn (New York: Springer).

Wang, Q., and Burkhalter, A. (2007). Area map of mouse visual cortex. J Comp Neurol 502, 339–357.

Wang, Q., Gao, E., and Burkhalter, A. (2007). In vivo transcranial imaging of connections in mouse visual cortex. J Neurosci Methods 159, 268–276.

Wang, Q., Gao, E., and Burkhalter, A. (2011). Gateways of ventral and dorsal streams in mouse visual cortex. J Neurosci 31, 1905–1918.

Wang, Q., Henry, A.M., Harris, J.A., Oh, S.W., Joines, K.M., Nyhus, J., Hirokawa, K.E., Dee, N., Mortrud, M., Parry, S., Ouellette, B., Caldejon, S., Bernard, A., Jones, A.R., Zeng, H., and Hohmann, J.G. (2014). Systematic comparison of adeno-associated virus and biotinylated dextran amine reveals equivalent sensitivity between tracers and novel projection targets in the mouse brain. J Comp Neurol 522, 1989–2012.

Wang, Q., Sporns, O., and Burkhalter, A. (2012). Network analysis of corticocortical connections reveals ventral and dorsal processing streams in mouse visual cortex. J Neurosci 32, 4386–4399.

Ypma, R.J., and Bullmore, E.T. (2016). Statistical Analysis of Tract-Tracing Experiments Demonstrates a Dense, Complex Cortical Network in the Mouse. PLoS Comput Biol 12, e1005104.

Zingg, B., Hintiryan, H., Gou, L., Song, M.Y., Bay, M., Bienkowski, M.S., Foster, N.N., Yamashita, S., Bowman, I., Toga, A.W., and Dong, H.W. (2014). Neural networks of the mouse neocortex. Cell 156, 1096–1111.

